# Modulation of endometrial E-Cadherin and N-Cadherin by ovarian steroids and embryonic stimuli

**DOI:** 10.1101/2021.04.08.438921

**Authors:** Abhishek Tiwari, Nancy Ashray, Neha Singh, Shipra Sharma, Deepak Modi

## Abstract

The endometrium is a dynamic tissue that undergoes extensive remodelling to attain a receptive state which is further modulated in presence of an embryo for successful initiation of pregnancy. Cadherins are the proteins of junctional complex of which E-cadherin (E-Cad) is crucial for maintaining epithelial cell state and integrity of the epithelial barrier; gain of N-cadherin (N-Cad) in epithelial cells leads to epithelial to mesenchymal transition (EMT). In the present study, we aimed to investigate the expression of E-Cad and N-Cad in the mouse endometrial luminal epithelium and its modulation by estrogen, progesterone and embryonic stimuli. We observed that E-Cad is diffusely expressed in the luminal epithelium of mouse endometrium during the estrus stage and upon estrogen treatment. It is apico-laterally and basolaterally sorted at the diestrus stage and in response to combined treatment of estrogen and progesterone. In 3D spheroids of human endometrial epithelial cells, combined treatment with estrogen and progesterone led to lateral sorting of E-Cad. In the mouse endometrium at the time of embryo implantation, there is loss of E-Cad which was associated with the gain of N-Cad suggestive of EMT in the luminal epithelium. This EMT is possibly driven by embryonic stimuli as treatment with estrogen and progesterone did not lead to gain of N-Cad expression. In conclusion, the present study demonstrates that steroid hormones directly affect E-Cad sorting in the endometrial epithelium.

## Introduction

Uterine endometrium consists of the simple columnar epithelium that extends in the supporting stroma to form the coiled endometrial glands. During the endometrial cycle, the stromal and epithelial cells undergo extensive remodeling under the influence of ovarian steroids; estrogen, and progesterone (Gellersen and Brosens, 2014; Deligdisch-Schor and Mareş Miceli, 2020). These modifications prepare the endometrium to endow receptivity for embryo apposition and implantation. Any defects in these processes can lead to implantation failure resulting in infertility (Evans *et al*., 2016).

Classical transplantation experiments suggest that the endometrium always rejects the embryo unless correctly primed by both estrogen and progesterone. However, if the luminal epithelium is scratched, the embryo readily implants even in the absence of hormonal priming (Ashary *et al*., 2018). These observations indicate that the integrity of the luminal epithelium is a determinant of the ability of the embryo to implant. In most mucosal surfaces, the integrity of the epithelium is maintained by junctional complexes which are composed of an array of proteins (Díaz-Díaz *et al*., 2020; Piprek *et al*., 2020). One key protein of the junctional complex is Epithelial-Cadherin (E-Cad) which is developmentally required for epithelial biogenesis (Gall and Frampton, 2013). In the uterus, E-Cad is essential for the biogenesis of epithelium, and mice knockout for *Cdh1* (gene encoding for E-Cad) have disorganized uterine epithelium (Reardon *et al*., 2012). Also during the menstrual cycle, when the endometrium undergoes cyclic remodeling, there are dynamic changes in the expression of E-Cad (Fujimoto *et al*., 1996; Matsuzaki *et al*., 2010). Further, the expression of E-Cad is altered in the endometrial epithelium of infertile women (Matsuzaki *et al*., 2010; Kakar-Bhanot *et al*., 2019; Sancakli Usta *et al*., 2020; Whitby *et al*., 2020) suggesting a key role of this protein in adult endometrium.

Beyond its role in the biogenesis of epithelium, E-Cad also plays a role in defining the apicobasal polarity of the epithelium and is a pre-requisite for directionality during the secretory and absorptive functions for maintenance of homeostasis (Desai *et al*., 2009; Gall and Frampton, 2013; Schnoor, 2015; Jiang *et al*., 2018; Piprek *et al*., 2020). Interestingly, there are stage-dependent changes in apico-lateral sorting of E-Cad in the endometrium. In the proliferative phase, E-Cad is predominantly diffused that gets laterally sorted in the secretory phase (Fujimoto *et al*., 1996). However, what regulates the apico-lateral sorting of E-Cad in the endometrial epithelium is yet not clear. The ovarian steroids; estrogen and progesterone are known to have profound effects on the endometrium and modulate the expression of many genes (Young, 2013; Evans *et al*., 2016; Deligdisch-Schor and Mareş Miceli, 2020). Whether the dynamic changes in E-Cad sorting in the endometrial epithelium is regulated by steroid hormones is unknown.

For a successful pregnancy, the embryo apposes on the receptive endometrium and implants in the endometrial bed. Extensive modifications occur in the endometrium at this time in response to the mid-cycle estrogen surge and embryonic factors (Nimbkar-Joshi *et al*., 2012; Zhang *et al*., 2013; Strug *et al*., 2016; Kurian and Modi, 2019; Mishra *et al*., 2021). In response to embryonic stimuli, the luminal epithelial cells lose their polarity, proliferate and there is an altered expression of several molecules required for embryo apposition and implantation (Nimbkar-Joshi *et al*., 2012; Modi and Bhartiya, 2015; Ashary *et al*., 2018; Laheri *et al*., 2018; Ochoa-Bernal and Fazleabas, 2020). Controversies exist on the pattern of E-Cad expression in the endometrial epithelium at the time of embryo implantation. Some studies have reported a loss of E-Cad in the endometrial epithelial cells at the site of embryo implantation, some have reported an increase in expression and/or higher apically sorted E-Cad while others have reported no change in expression of E-Cad at the time of implantation (Potter *et al*., 1996; Paria *et al*., 1999; Jha *et al*., 2006; Guo *et al*., 2009; Arora *et al*., 2016; Payan-Carreira *et al*., 2016; Dudley *et al*., 2017; Ashary *et al*., 2018; Yuan *et al*., 2018). Differences in fixatives used, staining methods, antibody epitopes could be possible reasons for such diverse observations.

For the embryo to implant, the luminal epithelium acts as a strong barrier that needs to be breached. Various mechanisms such as apoptosis, necroptosis and phagocytosis of epithelial cells by the trophoblasts have been proposed to aid in epithelial breaching (Li *et al*., 2015; Aplin and Ruane, 2017; Ashary *et al*., 2018; Akaeda *et al*., 2021). Beyond these, Epithelial to Mesenchymal Transition (EMT) is also proposed as a mechanism of embryo implantation. Using an *in vitro* model, it was shown that JAr spheroids (resembling early-stage human embryos) promote migration and expression of EMT markers in endometrial epithelial cells (Uchida *et al*., 2012; Ran *et al*., 2020). However, if luminal epithelial cells of the endometrium undergo EMT during embryo implantation *in vivo* is currently unknown. Further, if steroid hormones or embryonic signals mediate EMT in the endometrial luminal epithelium *in vivo* at the time of embryo implantation is not known.

In the present study, we aimed to investigate the effects of estrogen and progesterone on the dynamics of E-Cad sorting in the endometrial luminal epithelium at single-cell resolution. We also investigated the occurrence of EMT in the endometrial epithelium at the site of embryo implantation in the mouse.

## Methods

### Ethics Statement

The ethics committee at ICMR-NIRRH approved the use of animals under project number 01/16 and 18/17.

### Animal tissue collection

Uterine tissues were collected from C57BL/6 mice in estrus and diestrus stages (n=3 per stage) as determined by vaginal swabs. Females were allowed to mate with males of proven fertility and the day of vaginal plug was designated as day 1 of pregnancy. Animals were sacrificed on the noon of 3 days post-coitus (dpc), 4 dpc and 5 dpc (n=3 per dpc).

### Steroid hormone treatment

17β-estradiol (Cat No: E2758; Sigma-Aldrich, St. Louis, USA) and progesterone (Cat No: P8783; Sigma-Aldrich) were solubilized in ethanol and emulsified in corn oil (Sigma-Aldrich) overnight. Juvenile female mice (3 weeks old) were subcutaneously injected with either 1μg of estrogen or 1mg of progesterone or both for 5 consecutive days (n= 3 per treatment). The doses and schedule used in this study were as described elsewhere (Tibbetts *et al*., 1998; Galvankar *et al*., 2017; James *et al*., 2018). Controls were injected with an equivalent volume of ethanol in corn oil. On the 6^th^ day, (24h after the last dose) the uteri were collected and fixed overnight in 4% of paraformaldehyde (Sigma-Aldrich) and processed for paraffin embedding and sectioning.

### 3D Culture of human endometrial epithelial cells

Ishikawa cells are epithelial cells derived from the endometrium of a patient with adenocarcinoma and studies have shown that the Ishikawa cells are steroid hormone-responsive and mimic several aspects of the normal human endometrial epithelium (Daftary *et al*., 2002; Uchida *et al*., 2012; Laheri *et al*., 2018; Kakar-Bhanot *et al*., 2019). Ishikawa cells were purchased from American Type Culture Collection and cultured in DMEM/F12 supplemented with 10% (v/v) fetal bovine serum, 100 IU/ml penicillin, and 100 μg/ml streptomycin (all from Gibco, Life Technologies, Gaithersburg, MD, USA).

For the generation of spheroids, Ishikawa cells were trypsinized, washed, and mixed in 70% Matrigel (Gibco, Life Technologies, Gaithersburg, MD, USA) prepared in DMEM/F12 with 2% dextran charcoal-stripped FBS. Domes of ∼200μl (each containing ∼500 cells) were prepared on coverslips and cultured at 37^0^C in 5% CO2 with media change after every two days. After eight days, the spheroids were challenged with 10^−8^ M 17β-estradiol or 10^−6^ M progesterone alone or in combination for 24h. These concentrations and timings were chosen as they are known to alter the expression of several genes and proteins in Ishikawa cells (Daftary *et al*., 2002; Uchida *et al*., 2012; Laheri *et al*., 2018). The next day, the domes were washed in prewarmed phosphate-buffered saline (PBS) and fixed in 4% paraformaldehyde overnight.

### Immunofluorescence for tissue sections

5μm thick sections were deparaffinized in xylene, and rehydrated in decreasing grades of alcohol and immunofluorescence staining was carried out as described (Godbole *et al*., 2017; Shende *et al*., 2021). Antigen retrieval was done in 10 mM Tris EDTA buffer pH 9 at 95^0^C for 30 min, followed by incubation in freshly prepared 10 mM sodium borohydrate (Sigma-Aldrich) for 30 mins and washed in PBS. The sections were blocked in 5% bovine serum albumin (BSA) (MP Biochemicals Maharashtra, India, Cat No: 160069) followed by overnight incubation with anti-E-Cad (Abcam: ab15148; 1:50 dilution) or anti-N-Cad antibody (Abcam: ab18203; 1:200 dilution) prepared in PBS. This E-Cad antibody exclusively detects the extracellular domain allowing us to monitor the clustering of the functional region of E-Cad. Next day, the sections were washed three times in PBS, and the signal was detected with donkey anti-rabbit IgG Alexa Fluor 568 (Invitrogen: A10042; 1:1000 dilution). The sections were counterstained in 0.1 μg/ ml of DAPI (Sigma-Aldrich) and mounted in the ProLong Gold Antifade mountant (Cat No: P36930; Invitrogen,). For negative controls, the sections were incubated with PBS instead of primary antibody. Images were captured using an inverted fluorescence microscope equipped with an sCMOS camera (Leica Microsystems DMi8, Mannheim, Germany). The images were analyzed, and the quantification was done using Fiji version of ImageJ software.

### Immunofluorescence for spheroids

Overnight fixed spheroids in matrigel domes were washed extensively in PBS and blocking was done in 5% BSA for 1h in 6 well plates. The domes were incubated for 16-20h in E-Cad antibody (Abcam: ab15148; 1:50) at 4^0^C. Domes incubated with PBS were used as negative controls. The next day, domes were washed in PBS containing 0.1% Triton X-100 and incubated in Alexa Fluor 568 conjugated secondary donkey anti-rabbit antibody (Invitrogen: A10042; 1:1000) for 4h at room temperature. After washing in PBS with 0.1% TritonX-100, the domes nuclei were counter-stained in DAPI for 1h at room temperature and washed in PBS. The coverslips containing domes were gently mounted in ProLong Gold Antifade mountant and left overnight at 4^0^C. Next day, spheroids were imaged under a confocal microscope (Leica Microsystems DMi8, Mannheim, Germany). Spheroids with at least 15 cells were imaged and optical z-slices were acquired every 5-μm for a total of 500-μm. The numbers of cells with sorted E-Cad in each spheroid were manually counted by two independent observers.

### Quantification of E-Cad in single cells of tissue sections

For estimation of E-Cad signal intensities and sorting in the luminal epithelium, 5 independent areas of luminal epithelium were imaged. Lateral margins of individual cells were identified in grayscale images and fluorescence intensities from apical to basal end were quantified in ImageJ. Quantification was done on ten individual cells from five selected areas per animal (50 cells/animal). Data were collected from 3 individual replicates. 3D surface plots were generated for single cells using ImageJ.

To compare the distribution of E-Cad along the length of lateral membrane, intensity values of each pixel along the entire length of the individual cell were exported to Microsoft Excel to normalize for variation in cell height.

### Statistical analysis

The mean ± SD for all the experimental data was calculated and statistical analysis was done by Student’s t-test using GraphPad Prism, version 8. p < 0.05 was accepted as statistically significant.

## Results

### E-Cad is differentially sorted on lateral membranes of the luminal epithelial cells in estrus and diestrus stages

In luminal epithelial cells of the estrus stage endometrium, E-Cad was diffused throughout the cytoplasm with very limited staining on the lateral walls (Fig 1.A, B and C). In the diestrus stage, E-Cad mainly localized on to the lateral membranes with almost no staining in the cytoplasm (Fig 1.A, B and C). The negative controls did not show any staining (Supplementary Fig 1). In the estrus uteri, 3D surface plots revealed diffuse E-Cad staining throughout the cell (Fig 1.D). However, E-Cad distribution was highly polarized in luminal epithelium cells of the diestrus stage with two well-separated high-intensity peaks in the apical region. Also, there were two minor peaks observed at the basal end of the cells and their intensity was comparable to that in the estrus stage uterus (Fig 1.D).

**Figure 1.**
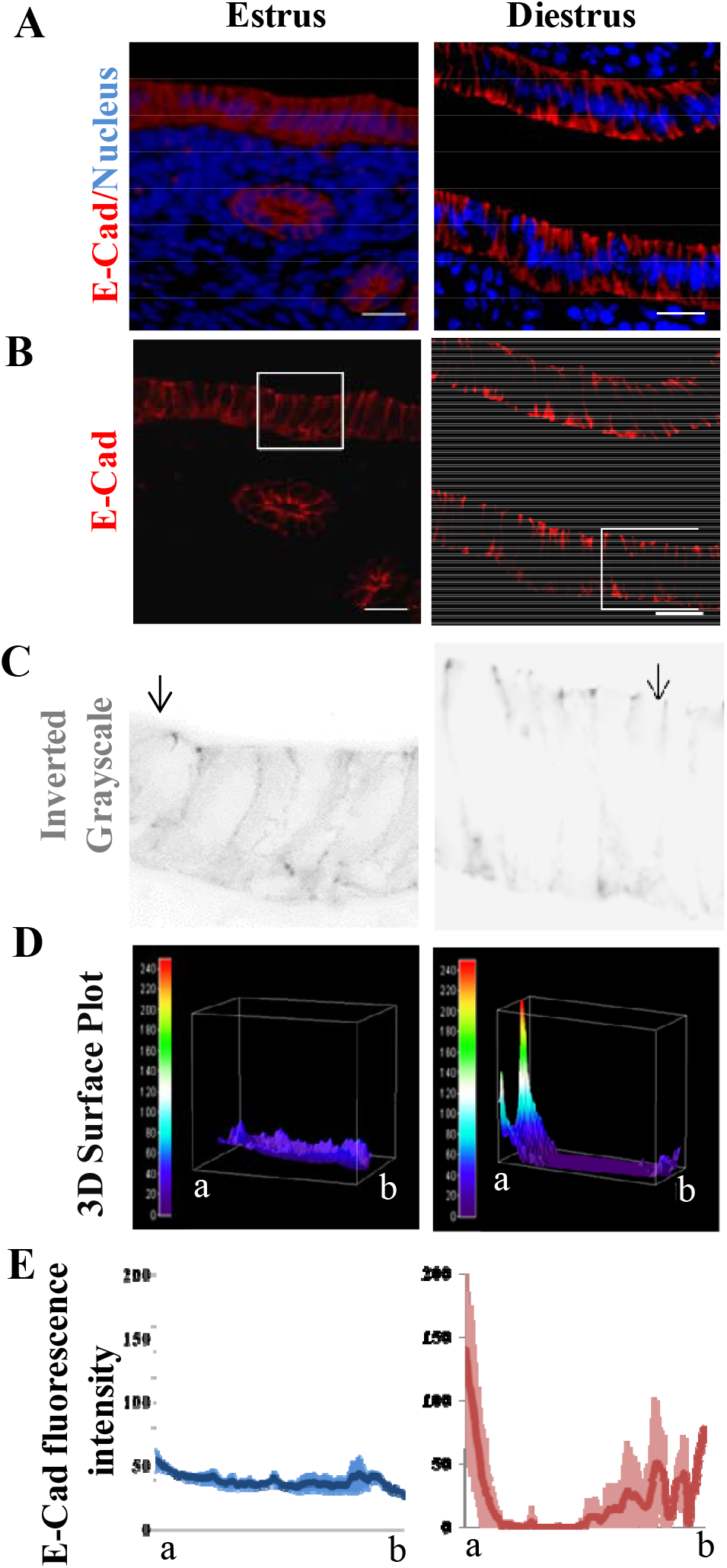
E-Cad is differentially sorted on the lateral membranes of the luminal epithelial cells of the endometrium in estrus and diestrus stages. (A) Immunofluorescence images for E-Cad (red) and nucleus (blue) at Estrus and Diestrus stages of mouse endometrium, Scale bar = 20µm. The negative control is shown in supplementary Fig 1 (B) Red channel images showing E-Cad staining along with selected area boxed for grayscale images. (C) Inverted grayscale images of the boxed area (in B) showing E-Cad sorting. Arrows indicate the apical end. (D) 3D surface plots of a single epithelial cell from each stage. Scale on the left represents the measure of thermal gradients. The apical end is denoted as “a” and basal end is denoted as “b”. (E) The intensity of E-Cad was measured on the lateral membranes of single cells of the luminal epithelium at estrus and diestrus. The X-axis is the distance from the apical to the basal end. Y-axis is the mean normalized fluorescence intensity of E-Cad. At each stage, data is mean of 150 single cells analyzed from 3 biological replicates. The shaded region denotes standard deviations

We quantified laterally distributed E-Cad in 150 individual cells of the luminal epithelium in estrus and diestrus stages (n=3/stage). At the estrus stage, E-Cad was minimally detected in the lateral membrane and was more or less uniformly distributed. At the diestrus stage, E-Cad was maximum apico-laterally followed by a steep decline in the middle region and a marginal increase towards the basal end (Fig 1.E).

### Steroid hormones modulate E-Cad sorting on the lateral membrane of endometrial luminal epithelium

To understand if ovarian steroids mediate E-Cad sorting, we analyzed its immunostaining in the endometrium of neonatal mice treated with estrogen or progesterone alone or in combination. In the luminal epithelial cells of controls, E-Cad was cytoplasmic and also found uniformly distributed on the lateral membranes (Fig 2 A, B, and C). In the luminal epithelial cells of the estrogen-treated mice E-Cad was diffused and was generally cytoplasmic with some apical staining (Fig 2 A, B and C). In response to progesterone treatment, E-Cad was sorted to the lateral membrane exclusively in the apical region (Fig 2 A, B and C). This was visualized as two prominent peaks at the apico-lateral side (Fig 2.D). Interestingly, in the endometria of mice treated with combined estrogen and progesterone, there was clear apico-lateral sorting of E-Cad along with the basal peaks similar to those observed in the diestrus stage endometrium (Fig 2.D). The negative controls did not show any staining (Supplementary Fig 1).

**Figure 2.**
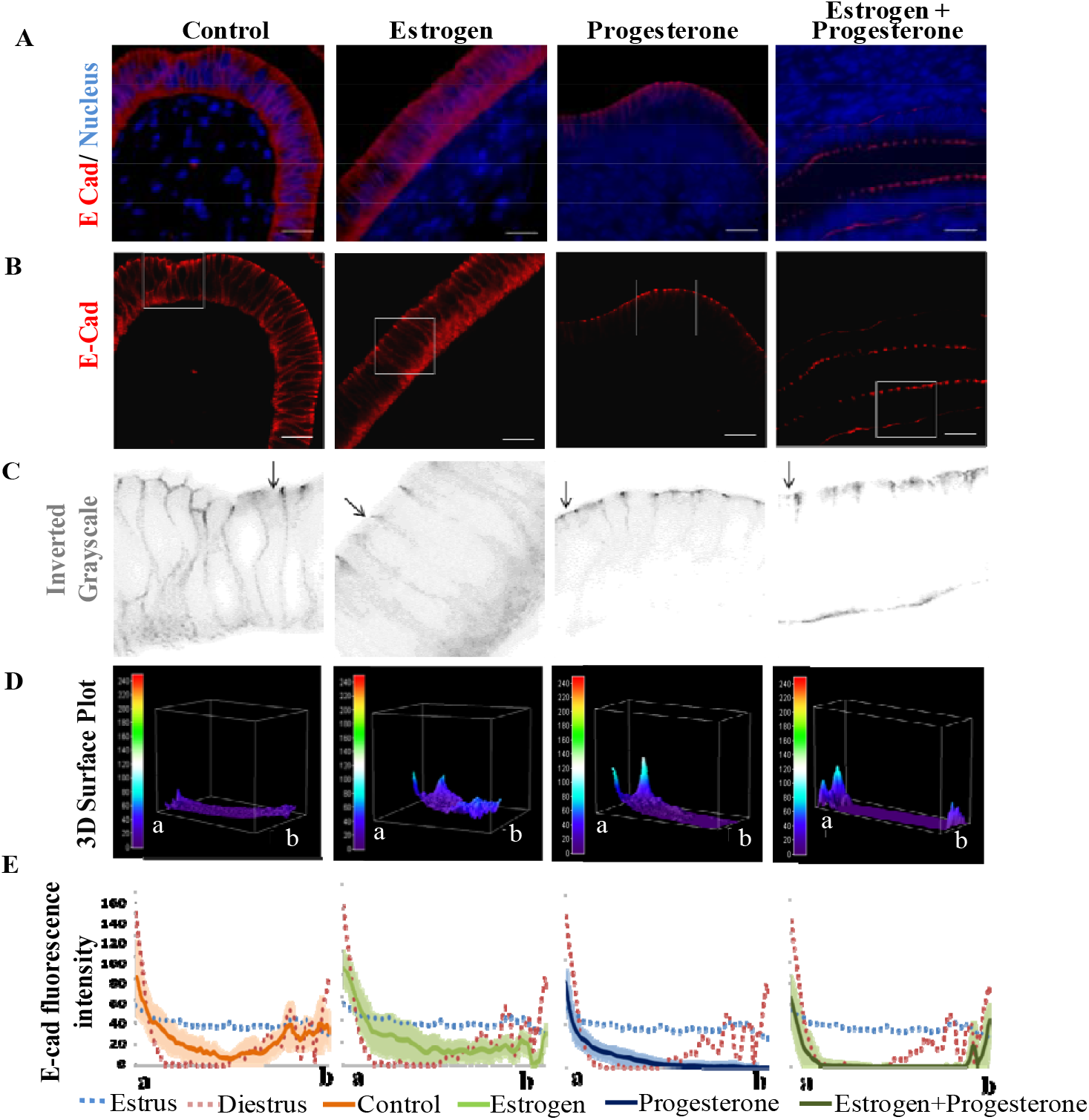
Steroid hormones modulate E-Cad sorting on the lateral membrane of endometrial luminal epithelium. Juvenile mice were treated with estrogen, progesterone alone or in combination for 5 days, and uteri sections were stained for E-Cad (A) Immunofluorescence images for E-Cad (red) and nucleus (blue) in the uteri treated with vehicle control, estrogen, progesterone, and estrogen + progesterone. The negative control is shown in Supplementary Fig 1. Scale bar = 20µm. Controls are animals at the same stage treated with vehicle. (B) Red channel images showing E-Cad staining along with selected area boxed for grayscale images. (C) Inverted grayscale images are obtained from the inset area in the red channel. Arrows indicate the apical end. (D) 3D surface plots for E-Cad intensity in a single epithelial cell. The scale on the left represents the measure of thermal gradients. The apical end is denoted as “a” and the basal end is denoted as “b”.(E) Intensity of E-Cad was measured on the lateral membranes of single cells of luminal epithelium in each treatment. The X-axis is the distance from the apical to basal end. Y-axis is the mean normalized fluorescence intensity of E-Cad. In each treatment, data is mean of 150 single cells analyzed from 3 biological replicates. The shaded region denotes standard deviations.

We quantified the intensity of E-Cad on lateral membranes of single cells (n=150) of the luminal epithelium of steroid-treated mice (n=3/group). In response to estrogen treatment, E-Cad was largely cytoplasmic with minimal but uniform distribution laterally (Fig 2.E). In mice treated with progesterone, most E-Cad was well sorted in the apico-lateral region with a steep decline posteriorly. The basal peaks of E-Cad as seen in the diestrus stage animals were not observed in response to progesterone treatment (Fig 2.E). In mice treated with both estrogen and progesterone, E-Cad was well sorted apico-laterally along with the basal peaks as observed in the diestrus stage endometrium.

### Steroid hormones modulate apico-lateral sorting of E-Cad in the lateral membrane of human endometrial epithelial cells

Human endometrial epithelial cells (Ishikawa) were cultured in 3D spheroids and challenged with estrogen or progesterone alone or in combination. In untreated control spheroids, E-Cad was mainly cytoplasmic in most cells. In estrogen-treated spheroids too, most E-Cad was cytoplasmic with a few cells having membrane-localized staining (Fig 3.A). Upon treatment with progesterone, E-Cad was detected on the membranes of cells with little cytoplasmic staining in most spheroids. In response to combined estrogen and progesterone treatment, intense membrane-localized staining for E-cad in the cells of most spheroids was observed (Fig 3.A). The negative control without primary antibody did not show any staining (Supplementary Fig 1).

**Figure 3.**
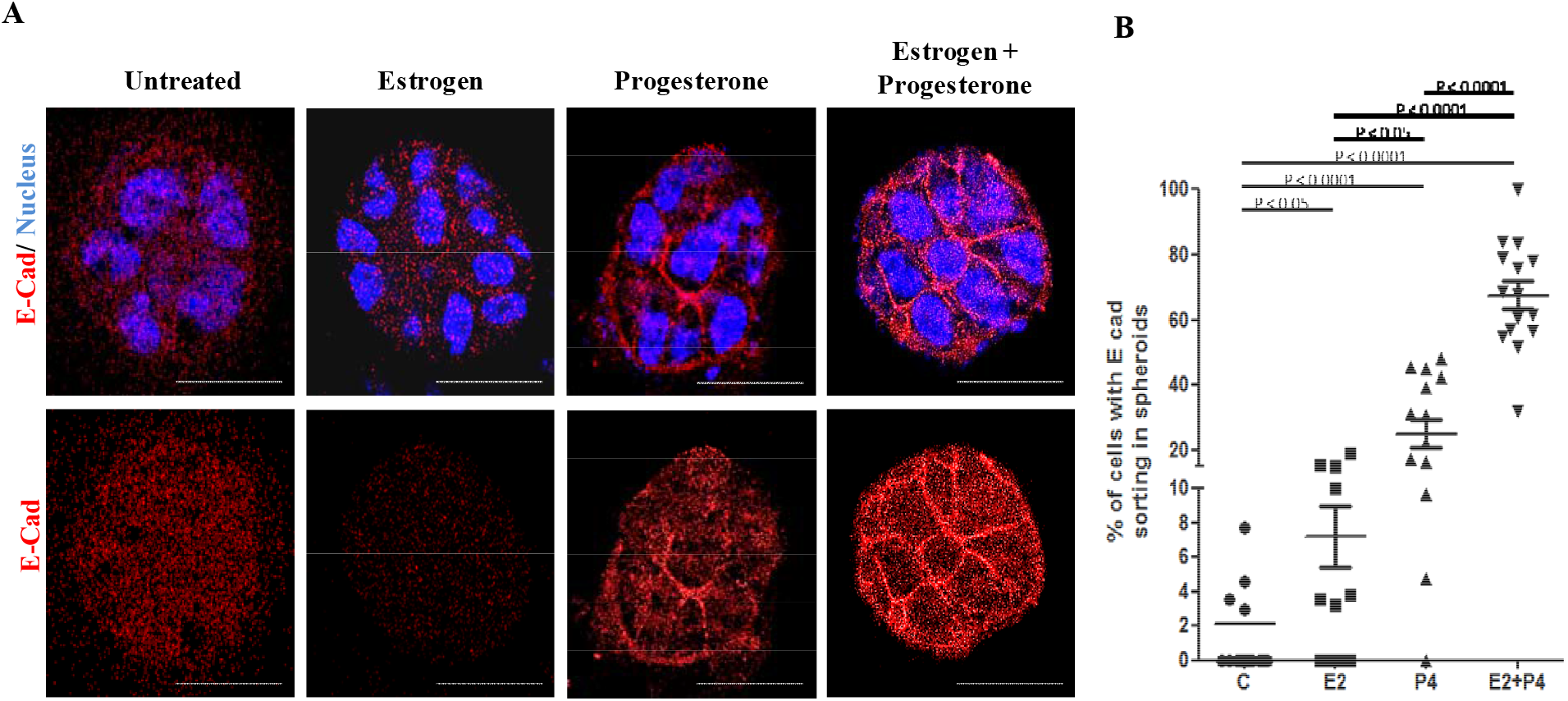
Steroids hormones modulate apico-lateral sorting of E-Cad in the lateral membrane of human endometrial epithelial cells. (A) Spheroids of human endometrial epithelial cells (Ishikawa cells) were challenged with estrogen or progesterone alone or in combination for 24hrs and stained for E-Cad by immunofluorescence. Controls are vehicle-treated spheroids. Each image represents the largest slice of the Z stack of an individual spheroid. Scale bar= 20µm. (B) Percentage of spheroids with membrane sorted E-Cad. Each dot represents a single spheroid (15-40 cells/spheroid) and 15 spheroids were counted per treatment condition. Mean ±SD values are shown. Statistically significant p values in comparison control and between treatment groups are given. The negative control incubated without a primary antibody is shown in Supplementary Fig 1.

We quantified the numbers of cells that showed membrane-localized E-Cad in individual spheroids (n=15/treatment group). In the untreated controls, ∼2% of cells/spheroid had sorted E-Cad, while in response to estrogen treatment almost 8-10% of cells had sorted E-Cad. In response to progesterone treatment, nearly 25% of cells had sorted E-Cad, almost 75% of cells had membrane sorted E-Cad in the combined estrogen and progesterone treatment group. This increase was statistically significant (p<0.0001) as compared to control and estrogen treatment (Fig 3.B).

### Expression of E-Cad in luminal epithelial cells at the time of embryo implantation

At 3 dpc, E-Cad was detected on the apico-lateral and baso-lateral sides of the cells of luminal epithelium similar to that seen in the diestrus stage endometrium (Fig 4.A, B and C); however, the intensity of the apical peak appeared reduced (Fig 4.D). On 4 dpc the intensity of E-Cad baso-laterally reduced as compared to 3 dpc. On 5 dpc, E-Cad was detected mainly towards the apico-lateral region, the baso-lateral staining was not observed (Fig 4.B, C and D). As compared to 3 and 4 dpc, the intensity of E-Cad apically also reduced, and the two peaks appeared much closer (Fig 4.D). This effect was specific to the site of implantation as 3D intensity profiles of E-Cad staining at the inter-implantation site (on 5 dpc) were similar to that observed in the epithelial cells of the day 3 stage endometrium (Fig 4.D). The negative control without the primary antibody did not show any staining (inset in Fig 4A)

**Figure 4.**
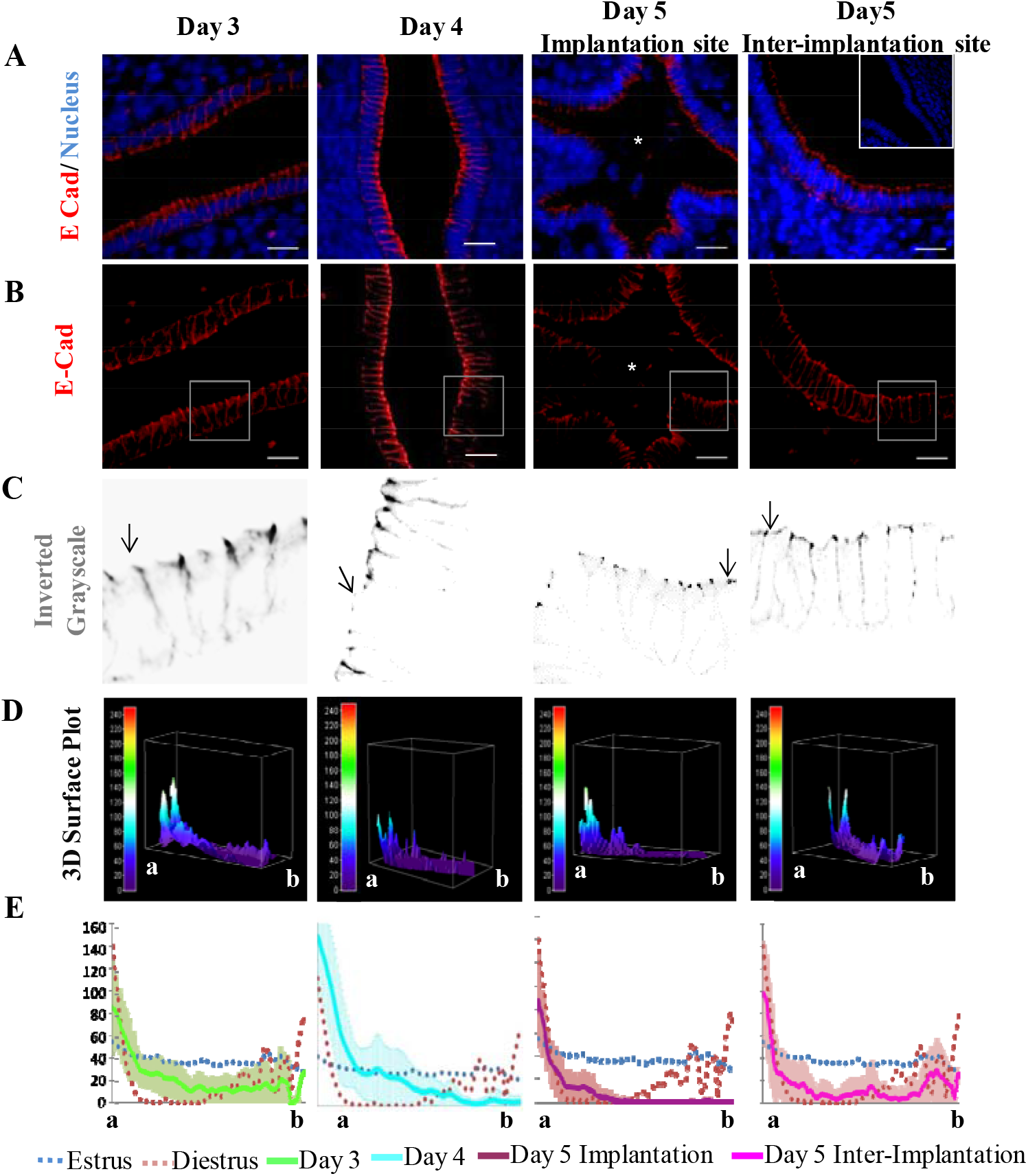
E-Cad is differentially sorted on the lateral membranes of the endometrial luminal epithelial cells during implantation under the influence of steroids. (A) Immunofluorescence images for E-Cad (red) and nucleus (blue) at Day 3 [3 days post coitus (dpc)], Day 4 (4 dpc), Day 5 implantation and inter-implantation site (5 dpc). * denotes the embryo. The negative control is shown in the inset. Scale bar=20µm (B) Red channel images showing E-Cad staining along with selected area boxed for grayscale images. (C) Inverted grayscale images of the boxed area showing E-Cad sorting. Arrows indicate apical end. (D) 3D surface plot of a single epithelial cell. The scale on the left represents the measure of thermal gradients. The apical end is denoted as “a” and the basal end is denoted as “b” (E) Intensity of E-Cad was measured on the lateral membranes of single cells of luminal epithelium at each stage. The X-axis is the distance from the apical to basal end. Y-axis is the mean normalized fluorescence intensity of E-Cad. At each stage, data is mean of 150 single cells analyzed from 3 biological replicates. The shaded region denotes standard deviations.

### EMT at the site of implantation

As compared to the preimplantation stages (3 dpc, n=3/stage), on 4 dpc when the endometrium is receptive and the embryo begins to appose, there was a significant (p<0.001) reduction in E-Cad expression in the epithelial cells (Fig 5.A). This reduction was largely due to the loss of E-Cad from the basolateral region. On 5 dpc, the embryo is in the implantation chamber, but the epithelial cell sloughing is yet not initiated. At this time, E-Cad was localized only apically in the epithelial cells. Quantitatively the levels of E-Cad on 5 dpc were significantly lower as compared to 3 dpc, the levels were comparable to those observed on 4 dpc (Fig 5.B). N-Cad was not detected in the luminal epithelium of the estrus, diestrus stage endometrium (not shown), it was also absent in the luminal epithelium on 3 and 4 dpc (Fig 5.A). On day 5 dpc, N-Cad staining was evident on the apical surface of the endometrial epithelium in the implantation chamber. The negative controls incubated without the primary antibody did not show any staining indicating the specificity of the reaction (Fig 5.A)

**Figure 5.**
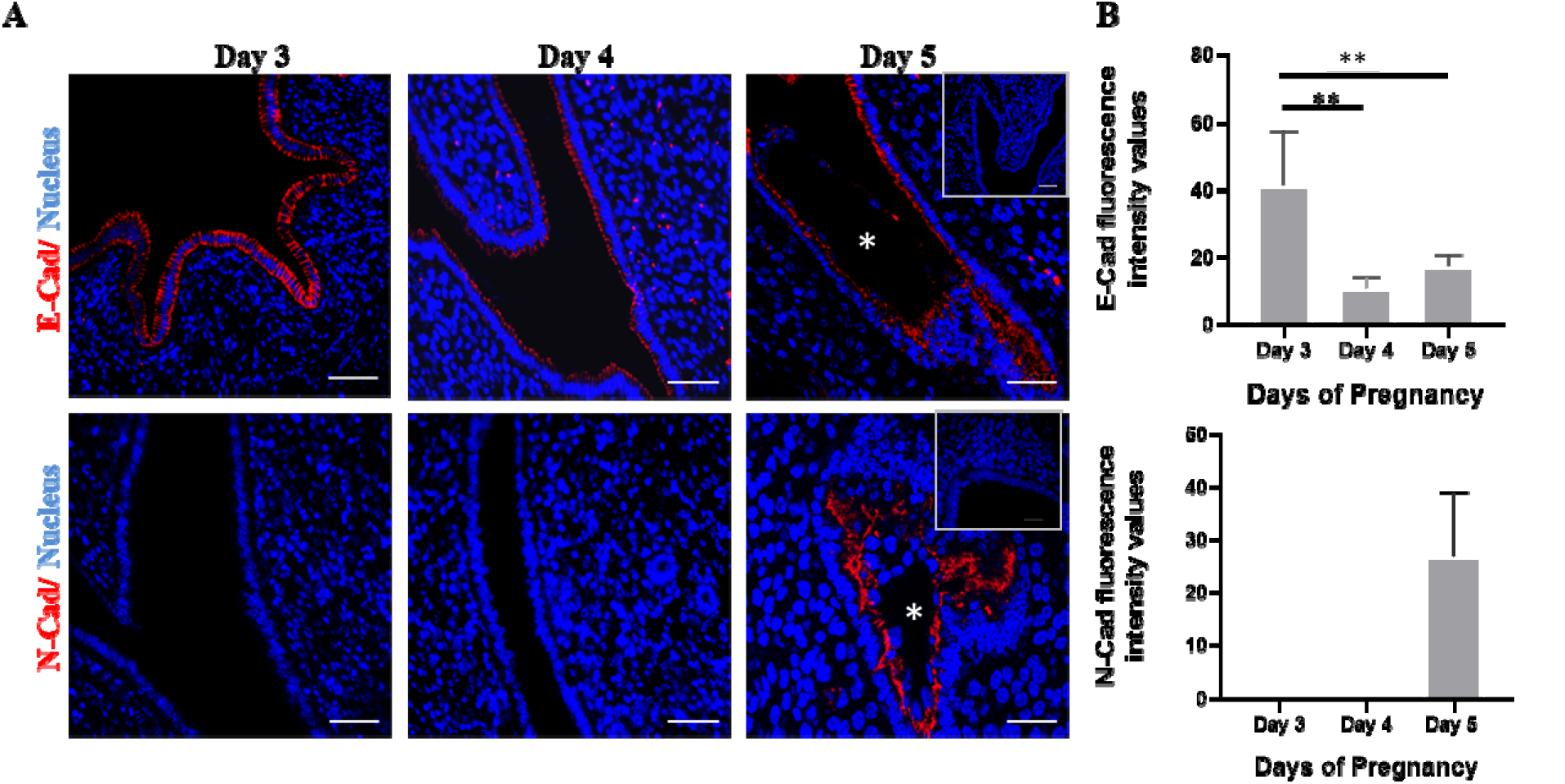
N-Cad expression is gained in the endometrial epithelium during the window of implantation. (A) Immunofluorescence images of E-Cad & N-Cad in early stages of pregnancy, i.e, Day 3 (3 dpc), Day 4 (4 dpc) & Day 5 (5 dpc). Scale bar = 50µm. The embryo is marked with an asterisk. The negative controls without the primary antibody are shown in the inset. (B) Fluorescence intensity of E-Cad/N-Cad in the endometrial epithelium during early days of pregnancy. Fluorescence values are the mean of 10 equal-sized areas from each technical replicate. n=3/each stage. Mean ± SD values are shown. Horizontal bars compare the groups and ** indicates statistical significance (p<0.05).

### Ovarian steroids do not induce N-Cad expression in the endometrial epithelium

To test if the gain in N-Cad expression in the endometrium on 5 dpc is in response to ovarian steroids, uterine sections of mice treated with estrogen and progesterone alone or in combination were obtained for N-Cad. No staining could be detected in any of the sections studied, albeit the positive control (mouse testis) was correctly stained for N-Cad (Fig 6).

**Figure 6.**
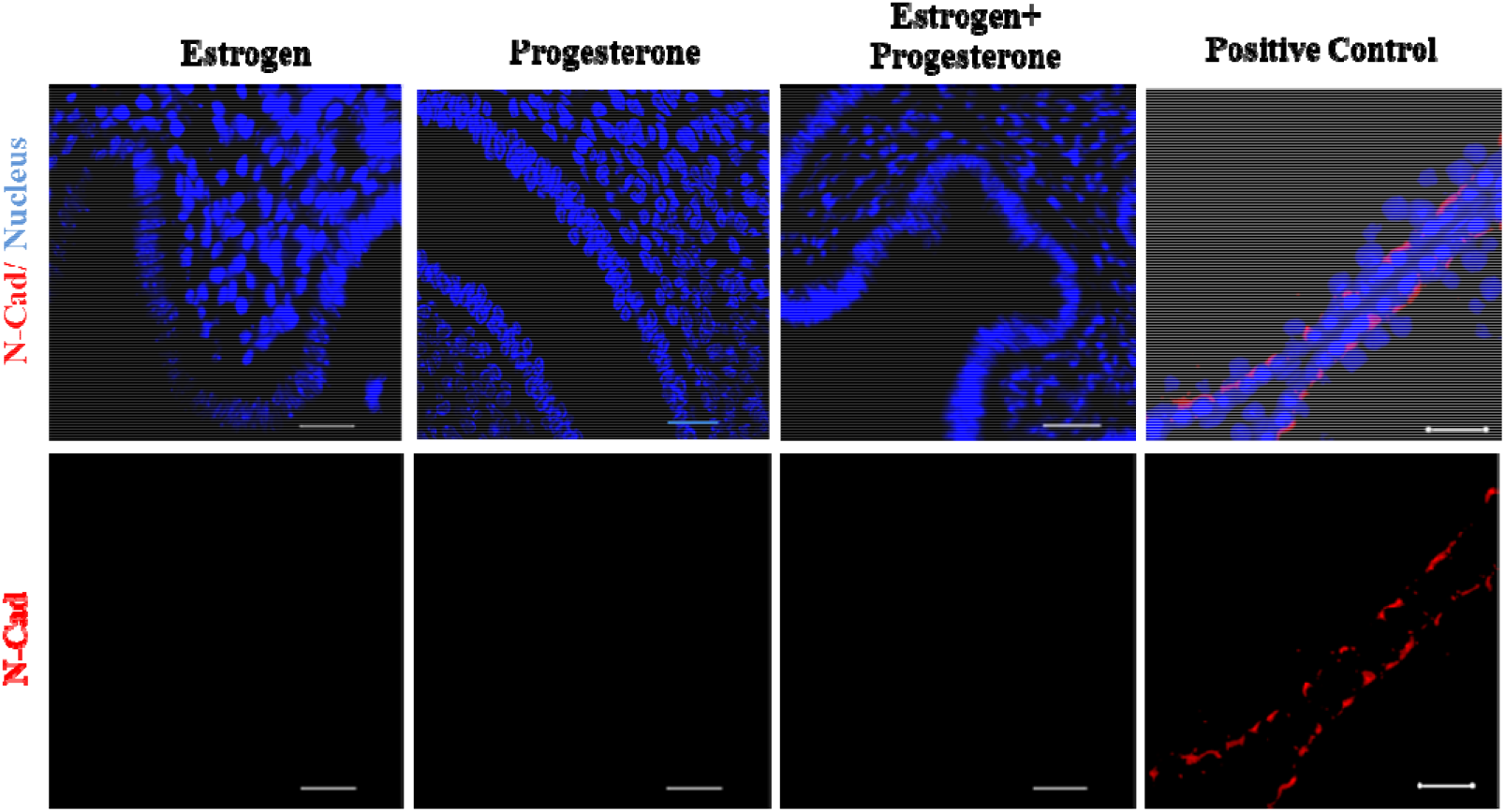
Steroid hormones do not modulate N-Cad expression in the endometrial epithelium in response to steroids. Juvenile mice were treated with estrogen, progesterone alone or in combination for 5 days and uteri sections were stained for N-Cad. The positive control is the adult mouse testis section. Scale bar = 20µm

## Discussion

In the present study, we demonstrate the differential sorting of E-Cad in mouse endometrial luminal epithelium during estrus and diestrus stages, and this sorting is modulated by estrogen and progesterone. We further show that at the time of implantation, expression of E-Cad is reduced and there is gain in N-Cad expression in the luminal epithelium of endometrium indicative of EMT.

E-Cad is a junctional protein that is required to maintain the apico-basal polarity, epithelial barrier integrity and tissue plasticity (Desai *et al*., 2009; Díaz-Díaz *et al*., 2020). This integrity and plasticity to an extent are achieved by a continuous turnover of E-Cad by cycles of endocytosis, sorting, and recycling back to the plasma membrane (Desai *et al*., 2009; Woichansky *et al*., 2016). Any alterations in these processes can have dramatic effects on tissue architecture and functions (Reardon *et al*., 2012; Gall and Frampton, 2013; Kakar-Bhanot *et al*., 2019; Whitby *et al*., 2020). Although the endometrial epithelium is thought to be columnar with anatomically distinct apical and basal ends, there are major differences in the distribution of adherence junction proteins in the proliferative and secretory stages of the human endometrium (Aplin and Ruane, 2017; Whitby *et al*., 2020). However, controversies exist on the distribution of E-Cad in epithelial cells during the endometrial cycle. Some studies have shown dynamic changes in E-Cad distribution in the proliferative and luteal phases (Fujimoto *et al*., 1996; Sancakli Usta *et al*., 2020). However, some studies have failed to confirm these findings and have shown no differences in E-Cad distribution in proliferative and secretory phases (Poncelet *et al*., 2002). In the present study, using an antibody that detects the extracellular domain, we observed that E-Cad is differentially distributed between the estrus and diestrus stages of mouse endometrium. At the estrus stage, E-Cad is largely cytoplasmic with minimal staining on the lateral membrane. In the diestrus stage endometrium, E-Cad is exclusively sorted in the apico-lateral and basolateral borders of luminal epithelial cells with negligible staining in the middle region. Interestingly, in the estrus stage, epithelial cells are loosely associated with each other while in the diestrus stage the epithelial cells are tightly bound laterally (Potter *et al*., 1996). These results indicate that the endometrial epithelium has dynamic changes in its plasticity and integrity during the different stages of the cycle and this is potentially due to altered sorting of E-Cad. The sorting of E-Cad in endometrial epithelium appears to be crucial, as reduced levels of laterally sorted E-Cad are observed in receptive phase endometrium of women with unexplained infertility (Kakar-Bhanot *et al*., 2019).

The dynamic remodeling of the endometrium that occurs during the estrus cycle is tightly regulated by ovarian steroid estrogen and progesterone. Our results show that the change in the distribution of E-Cad during estrus cycle is regulated by ovarian steroids. In the estrus stage, E-Cad was mainly cytoplasmic and not sorted apico-laterally. Analogous to this, estrogen treatment was associated with diffused cytoplasmic E-Cad in the epithelial cells of mouse endometrium and 3D spheroids of human endometrial epithelial cells. In a previous study, it was shown that estrogen aids in the cleavage of the extracellular domain of E-Cad in mouse epithelial cells (Potter *et al*., 1996). Since the antibody used in this study recognizes the extracellular domain of E-Cad, it is tempting to propose that the loss of E-Cad observed on the lateral walls in the estrus phase and in response to estrogen could be due to the cleavage of E-Cad extracellular domain. These results imply that the cyclic disruption of uterine epithelial integrity is dependent on the modification of E-Cad including its extracellular domain cleavage mediated by estrogen.

Like that observed in the diestrus stage endometrium, progesterone treatment led to localization of E-Cad apically on the lateral membranes with minimal cytoplasmic staining. Corroborating this, progesterone treated 3D spheroids of human endometrial epithelium also had E-Cad sorted to the lateral membrane. This is possibly a direct effect of the hormones on epithelial cells as the pattern could be recapitulated in the spheroids that are devoid of the endometrial stroma. We observed that while progesterone is necessary to localize E-Cad to lateral membranes, it was not sufficient to fully recapitulate the E-Cad distribution as seen in the diestrus stage. In the diestrus stage endometrium, the E-Cad was maximal in the apico-lateral membrane followed by a steep decline and then an increase at the basolateral side. This staining pattern was faithfully recapitulated in the luminal epithelial cells of mice treated with progesterone after estrogen priming and in 3D spheroids of human endometrial epithelial cells treated with both estrogen and progesterone. Thus unlike in the case of the estrus stage where estrogen promotes loss of E-Cad, in presence of progesterone, estrogen perhaps aids in basolateral sorting of E-Cad. In the endometrial epithelium, the GTPases Rab11 sorts E-Cad to apico-lateral regions (Kakar-Bhanot *et al*., 2019). Interestingly, both estrogen and progesterone regulate the expression of Rab11 and the Rab11 regulator, Rab coupling protein (Chen *et al*., 1999; Patil *et al*., 2005). Thus we propose that the cycle-dependent changes in the localization of E-Cad on the endometrial lateral membrane could be due to the effects of the steroid hormones on E-Cad extracellular cleavage and their effects on transcription of Rab11 and its partner proteins.

In mouse luminal epithelium, there is a dramatic cell loss following normal estrus and if pregnancy ensues, this cell loss is averted during the first 3 days till embryo invasion is initiated (Wang and Dey, 2006). Indeed, we observed that on day 3 when the endometrial epithelium is intact, E-Cad was distributed through the lateral walls of the luminal epithelial cells similar to that observed in diestrus endometrium in the non-pregnant state. Interestingly, we found that almost a day before the beginning of embryo invasion when the epithelial cell layer is intact (4 dpc), E-Cad was reduced from the basolateral membrane of luminal epithelial cells, the apical staining was however maintained. Further on day 5 (5 dpc), when embryo is in implantation chamber, E-Cad expression was strong apico-laterally and basal expression was not observed. This effect was specific to the sites of implantation as at the inter-implantation sites, distribution of E-Cad in luminal epithelial cells was identical to that of the diestrus stage. Loss of E-Cad from lateral membrane is observed in the canine, mouse and marsupial endometrium at the time of embryo implantation (Jha *et al*., 2006; Guo *et al*., 2009; Arora *et al*., 2016; Dudley *et al*., 2017; Ashary *et al*., 2018; Yuan *et al*., 2018; Bhurke *et al*., 2020). Interestingly, in species with fusion type of implantation (like the rabbit) which does not involve epithelial cell sloughing, high expression of E-Cad is detected in the basal plasma membrane at the time of embryo implantation (Olson *et al*., 1998). These observations imply that the luminal epithelium prepares itself by loosening the junctional complexes long before the actual process of invasion has begun. Although it is proposed that loss of lateral E-Cad is due to proteolytic cleavage of protein in response to estrogen surge at the time of implantation (Potter *et al*., 1996), it is important to note that the loss of E-Cad from lateral membrane occurs exclusively at the site of implantation, with patterns similar to diestrus stage endometrium in the inter-implantation sites. Furthermore, there was an intense accumulation of E-Cad at apico-lateral ends of luminal epithelial cells at the implantation sites. A similar apical accumulation of several integrins is observed in the monkey endometrium at the time of embryo implantation (Aplin and Ruane, 2017). Thus, it is likely that the fluidity of the membranes of luminal epithelial cells may change during implantation leading to the accumulation of E-Cad and other proteins on the apical portion of epithelium. However, the functional significance of such apically sorted E-Cad during embryo implantation needs to be investigated.

For embryo implantation, breaching of the epithelial barrier is thought to occur by removal of the cells by apoptosis, or phagocytosis of luminal epithelial cells by the embryonic trophoblasts (Ashary *et al*., 2018; Akaeda *et al*., 2021). Beyond these, using an i*n vitro* model, it has been shown that JAr spheroids alter E-Cad expression and induce EMT in endometrial epithelial cells (Uchida *et al*., 2012; Ran *et al*., 2020; Stern-Tal *et al*., 2020). Interestingly, we observed that by noon of 5 dpc, just before the beginning of embryo invasion, there is a significant loss of E-Cad and a gain of N-Cad in luminal epithelial cells suggestive of EMT. An earlier study has shown that EMT occurs in the mouse uterus during decidualization and this is regulated by EMT-specific transcription factor *Twist2* (Gou *et al*., 2019). However, in that study, EMT was studied in total tissue lysates and the cell types that undergo EMT is unknown. To the best of our knowledge, this is the first study demonstrating the occurrence of EMT in the luminal epithelium of endometrium during embryo implantation *in vivo*. It will be of interest to dissect the time kinetics and other markers of EMT in the endometrium to understand its involvement in breaching of epithelia for embryo invasion.

What would induce EMT in the luminal epithelium of the endometrium is a matter of investigation. An earlier study has shown that EMT could be induced in Ishikawa cells with the treatment of estrogen and progesterone even in absence of embryonic stimuli (Uchida *et al*., 2012). However, we did not detect any gain in expression of N-Cad in the luminal epithelial cells of endometrium obtained from mice treated with estrogen and progesterone alone or in combination. Thus it appears that *in vivo*, there could be additional factors required to induce EMT at the time of embryo invasion. Since embryo-derived factors alter the expression of genes in the luminal epithelium (Nimbkar-Joshi *et al*., 2012; Ashary *et al*., 2018, 2020; Laheri *et al*., 2018), it is tempting to propose the EMT occurring in luminal epithelium at the time of implantation could be due to embryo driven factors. Further studies will be required to address this hypothesis.

To summarize, in the present study we demonstrate that steroid hormones directly affect E-Cad sorting in the endometrial epithelium. This dynamic sorting would be required for the constant remodeling of endometrial epithelium during the cycle and also at implantation. Embryonic factors would drive EMT in the endometrial luminal epithelium at the time of embryo implantation. It would be interesting to study the downstream signaling of steroid hormones and other embryonic factors to understand the mechanism of E-Cad sorting and EMT in the endometrium for a successful pregnancy. This knowledge would help us in constructing organoids for studying human implantation and fertility-related disorders. In a long run, it will aid in understanding the causes of infertility and the management of recurrent implantation failure.

## Acknowledgments

We are grateful to Dr. Geetanjali Sachdeva (ICMR-NIRRH) for kindly sharing the Ishikawa cells. DM lab is funded by grants from the Indian Council of Medical Research (ICMR), Govt. of India. NA is thankful to DST-SERB for the project fellowship. NS is thankful to DBT project. NS and NA are grateful to ICMR for Seniro research fellowship. The project was funded by grants from DST-SERB to DM (DST No: CRG/2018/002314). The manuscript bears the NIRRH ID: RA/1055/04-2021.

---

**Supplementary Fig 1.**
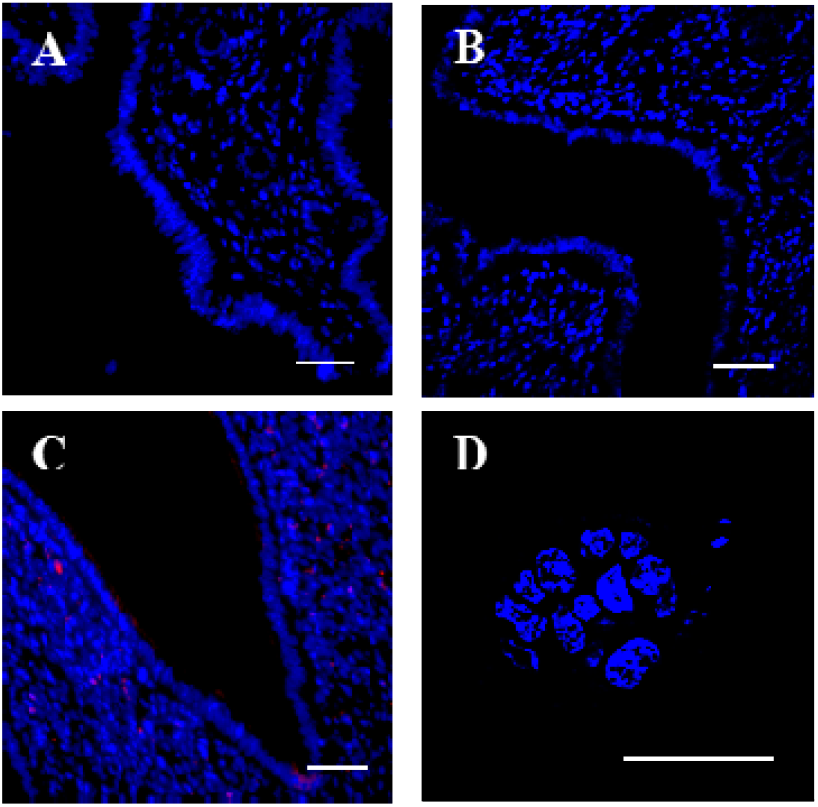
Negative controls for immunohistochemistry. All immunohistochemical stainings were accompanied by a negative control where the primary antibody was replaced by PBS. A) Estrus stage uterus, B) Diestrus stage uterus, C) Uterus on 4 dpc and D) Ishikawa cell spheroid treated with estrogen and progesterone. In A, B, and C Scale Bar=50µm. In D Scale bar is 20µm. Blue are DAPI stained nuclei.

## References

Akaeda S, Hirota Y, Fukui Y, Aikawa S, Shimizu□Hirota R, Kaku T, Gebril M, Hirata T, Hiraoka T, Matsuo M et al. (2021) Retinoblastoma protein promotes uterine epithelial cell cycle arrest and necroptosis for embryo invasion. EMBO Reports 22 e50927.

Aplin JD and Ruane PT (2017) Embryo-epithelium interactions during implantation at a glance. Journal of Cell Science 130 15–22.

Arora R, Fries A, Oelerich K, Marchuk K, Sabeur K, Giudice LC and Laird DJ (2016) Insights from imaging the implanting embryo and the uterine environment in three dimensions. Development 143 4749–4754.

Ashary N, Tiwari A and Modi D (2018) Embryo Implantation: War in times of love. Endocrinology 159 1188–1198.

Ashary N, Laheri S and Modi D (2020) Homeobox genes in endometrium: from development to decidualization. The International Journal of Developmental Biology 64 227–237.

Bhurke A, Kannan A, Neff A, Ma Q, Laws MJ, Taylor RN, Bagchi MK and Bagchi IC (2020) A hypoxia-induced Rab pathway regulates embryo implantation by controlled trafficking of secretory granules. Proceedings of the National Academy of Sciences of the United States of America 117 14532–14542.

Chen D, Ganapathy P, Zhu LJ, Xu E, Li Q, Bagchi IC and Bagchi MK (1999) Potential regulation of membrane trafficking by estrogen receptor α via induction of rab11 in uterine glands during implantation. Molecular Endocrinology 13 993–1004.

Daftary GS, Troy PJ, Bagot CN, Young SL and Taylor HS (2002) Direct regulation of beta3-integrin subunit gene expression by HOXA10 in endometrial cells. Molecular Endocrinology 16 571–579.

Deligdisch-Schor L and Mareş Miceli A (2020) Hormonal biophysiology of the uterus. In Advances in Experimental Medicine and Biology, pp 1–12. Springer.

Desai RA, Gao L, Raghavan S, Liu WF and Chen CS (2009) Cell polarity triggered by cell-cell adhesion via E-cadherin. Journal of Cell Science 122 905–911.

Díaz-Díaz C, Baonza G and Martín-Belmonte F (2020) The vertebrate epithelial apical junctional complex: Dynamic interplay between Rho GTPase activity and cell polarization processes. Biochimica et Biophysica Acta - Biomembranes 1862 183398.

Dudley JS, Murphy CR, Thompson MB and McAllan BM (2017) Epithelial cadherin disassociates from the lateral plasma membrane of uterine epithelial cells throughout pregnancy in a marsupial. Journal of Anatomy 231 359–365.

Evans J, Salamonsen LA, Winship A, Menkhorst E, Nie G, Gargett CE and Dimitriadis E (2016) Fertile ground: Human endometrial programming and lessons in health and disease. Nature Reviews Endocrinology 12 654–667.

Fujimoto J, Ichigo S, Hori M and Tamaya T (1996) Alteration of E-cadherin, α- and β-catenin mRNA expression in human uterine endometrium during the menstrual cycle. Gynecological Endocrinology 10 187–191.

Gall TMH and Frampton AE (2013) Gene of the month: E-cadherin (CDH1). Journal of Clinical Pathology 66 928–932.

Galvankar M, Singh N and Modi D (2017) Estrogen is essential but not sufficient to induce endometriosis. Journal of Biosciences 42 251–263.

Gellersen B and Brosens JJ (2014) Cyclic decidualization of the human endometrium in reproductive health and failure. Endocrine Reviews 35 851–905.

Godbole G, Suman P, Malik A, Galvankar M, Joshi N, Fazleabas A, Gupta SKSK and Modi D (2017) Decrease in Expression of HOXA10 in the Decidua After Embryo Implantation Promotes Trophoblast Invasion. Endocrinology 158 2618–2633.

Gou J, Hu T, Li L, Xue L, Zhao X, Yi T and Li Z (2019) Role of epithelial-mesenchymal transition regulated by twist basic helix-loop-helix transcription factor 2 (Twist2) in embryo implantation in mice. Reproduction, Fertility and Development 31 932–940.

Guo B, Tian Z, Han BC, Zhang XM, Yang ZM and Yue ZP (2009) Expression and Hormonal Regulation of Hoxa10 in Canine Uterus during the Peri-implantation Period. Reproduction in Domestic Animals.

James K, Bhartiya D, Ganguly R, Kaushik A, Gala K, Singh P and Metkari SM (2018) Gonadotropin and steroid hormones regulate pluripotent very small embryonic-like stem cells in adult mouse uterine endometrium. Journal of Ovarian Research 11 1–20.

Jha RK, Titus S, Saxena D, Kumar PG and Laloraya M (2006) Profiling of E-cadherin, β-catenin and Ca2+ in embryo-uterine interactions at implantation. FEBS Letters 580 5653–5660.

Jiang H, Shen J and Ran Z (2018) Epithelial-mesenchymal transition in Crohn’s disease. Mucosal Immunology 11 294–303.

Kakar-Bhanot R, Brahmbhatt K, Chauhan B, Katkam RR, Bashir T, Gawde H, Mayadeo N, Chaudhari UK and Sachdeva G (2019) Rab11a drives adhesion molecules to the surface of endometrial epithelial cells. Human Reproduction 34 519–529.

Kurian NK and Modi D (2019) Extracellular vesicle mediated embryo-endometrial cross talk during implantation and in pregnancy. Journal of Assisted Reproduction and Genetics 36 189–198.

Laheri S, Ashary N, Bhatt P and Modi D (2018) Oviductal glycoprotein 1 (OVGP1) is expressed by endometrial epithelium that regulates receptivity and trophoblast adhesion. Journal of Assisted Reproduction and Genetics 35 1419–1429.

Li Y, Sun X and Dey SK (2015) Entosis allows timely elimination of the luminal epithelial barrier for embryo implantation. Cell Reports 11 358–365.

Matsuzaki S, Darcha C, Maleysson E, Canis M and Mage G (2010) Impaired Down-Regulation of E-Cadherin and β-Catenin Protein Expression in Endometrial Epithelial Cells in the Mid-Secretory Endometrium of Infertile Patients with Endometriosis. The Journal of Clinical Endocrinology & Metabolism 95 3437–3445.

Mishra A, Ashary N, Sharma R and Modi D (2021) Extracellular vesicles in embryo implantation and disorders of the endometrium. American Journal of Reproductive Immunology 85 e13360.

Modi DN and Bhartiya P (2015) Physiology of embryo-endometrial cross talk. Biomed Res J 2 83–104.

Nimbkar-Joshi S, Katkam RR, Chaudhari UK, Jacob S, Manjramkar DD, Metkari SM, Hinduja I, Mangoli V, Desai S, Kholkute SD et al. (2012) Endometrial epithelial cell modifications in response to embryonic signals in bonnet monkeys (Macaca radiata). Histochemistry and Cell Biology 138 289–304.

Ochoa-Bernal MA and Fazleabas AT (2020) Physiologic events of embryo implantation and decidualization in human and non-human primates. International Journal of Molecular Sciences 21 1973.

Olson GE, Winfrey VP, Matrisian PE, NagDas SK and Hoffman LH (1998) Blastocyst-dependent upregulation of metalloproteinase/disintegrin MDC9 expression in rabbit endometrium. Cell and Tissue Research 293 489–498.

Paria BC, Zhao X, Das SK, Dey SK and Yoshinaga K (1999) Zonula occludens-1 and E-cadherin are coordinately expressed in the mouse uterus with the initiation of implantation and decidualization. Developmental Biology 208 488–501.

Patil VSS, Sachdeva G, Modi DNN, Katkam RRR, Manjramkar DDD, Hinduja I and Puri CPP (2005) Rab coupling protein (RCP): A novel target of progesterone action in primate endometrium. Journal of Molecular Endocrinology 35 357–372.

Payan-Carreira R, Pires MA, Santos C, Holst BS, Colaço J and Rodriguez-Martinez H (2016) Immunolocalization of E-cadherin and β-catenin in the cyclic and early pregnant canine endometrium. Theriogenology 86 1092–1101.

Piprek RP, Kloc M, Mizia P and Kubiak JZ (2020) The central role of cadherins in gonad development, reproduction, and fertility. International Journal of Molecular Sciences 21 1–21.

Poncelet C, Leblanc M, Walker-Combrouze F, Soriano D, Feldmann G, Madelenat P, Scoazec J-Y and Daraï E (2002) Expression of cadherins and CD44 isoforms in human endometrium and peritoneal endometriosis. Acta Obstetricia et Gynecologica Scandinavica 81 195–203.

Potter SW, Gaza G and Morris JE (1996) Estradiol induces E-cadherin degradation in mouse uterine epithelium during the estrous cycle and early pregnancy. Journal of Cellular Physiology 169 1–14.

Ran J, Yang HH, Huang HP, Huang HL, Xu Z, Zhang W and Wang ZX (2020) ZEB1 modulates endometrial receptivity through epithelial-mesenchymal transition in endometrial epithelial cells in vitro. Biochemical and Biophysical Research Communications 525 699–705.

Reardon SN, King ML, II JAML, Mann JL, DeMayo FJ, Lydon JP and Hayashi K (2012) Cdh1 is essential for endometrial differentiation, gland development, and adult function in the mouse uterus. Biology of Reproduction 86 1–10.

Sancakli Usta C, Turan G, Bulbul CB, Usta A and Adali E (2020) Differential expression of Oct-4, CD44, and E-cadherin in eutopic and ectopic endometrium in ovarian endometriomas and their correlations with clinicopathological variables. Reproductive Biology and Endocrinology 18 116.

Schnoor M (2015) E-cadherin Is Important for the Maintenance of Intestinal Epithelial Homeostasis Under Basal and Inflammatory Conditions. Digestive Diseases and Sciences 60 816–818.

Shende P, Gaikwad P, Gandhewar M, Ukey P, Bhide A, Patel V, Bhagat S, Bhor V, Mahale S, Gajbhiye R et al. (2021) Persistence of SARS-CoV-2 in the first trimester placenta leading to transplacental transmission and fetal demise from an asymptomatic mother. Human Reproduction 36 899–906.

Stern-Tal D, Achache H, Jacobs Catane L, Reich R and Tavor Re’em T (2020) Novel 3D embryo implantation model within macroporous alginate scaffolds. Journal of Biological Engineering 14 18.

Strug MR, Su R, Young JE, Dodds WG, Shavell VI, Dyíaz-Gimeno P, Ruyíz-Alonso M, Simón C, Lessey BA, Leach RE et al. (2016) Intrauterine human chorionic gonadotropin infusion in oocyte donors promotes endometrial synchrony and induction of early decidual markers for stromal survival: A randomized clinical trial. Human Reproduction 31 1552–1561.

Tibbetts TA, Mendoza-Meneses M, O’Malley BW and Conneely OM (1998) Mutual and intercompartmental regulation of estrogen receptor and progesterone receptor expression in the mouse uterus. Biology of Reproduction 59 1143–1152.

Uchida H, Maruyama T, Nishikawa-Uchida S, Oda H, Miyazaki K, Yamasaki A and Yoshimura Y (2012) Studies using an in vitro model show evidence of involvement of epithelial-mesenchymal transition of human endometrial epithelial cells in human embryo implantation. Journal of Biological Chemistry 287 4441–4450.

Wang H and Dey SK (2006) Roadmap to embryo implantation: clues from mouse models. Nature Reviews Genetics 7 185–199.

Whitby S, Zhou W and Dimitriadis E (2020) Alterations in Epithelial Cell Polarity During Endometrial Receptivity: A Systematic Review. Frontiers in Endocrinology 11.

Woichansky I, Beretta CA, Berns N and Riechmann V (2016) Three mechanisms control E-cadherin localization to the zonula adherens. Nature Communications 7 1–11.

Young SL (2013) Oestrogen and progesterone action on endometrium: A translational approach to understanding endometrial receptivity. Reproductive BioMedicine Online 27 497–505.

Yuan J, Deng W, Cha J, Sun X, Borg JP and Dey SK (2018) Tridimensional visualization reveals direct communication between the embryo and glands critical for implantation. Nature Communications 9 1–13.

Zhang S, Lin H, Kong S, Wang S, Wang H, Wang H and Armant DR (2013) Physiological and molecular determinants of embryo implantation. Molecular Aspects of Medicine 34 939–980.

